# Gene-rich X chromosomes implicate intragenomic conflict in the evolution of bizarre genetic systems

**DOI:** 10.1101/2020.10.04.325340

**Authors:** Noelle Anderson, Kamil S. Jaron, Christina N. Hodson, Matthew B. Couger, Jan Ševčík, Brooke Weinstein, Stacy Pirro, Laura Ross, Scott William Roy

## Abstract

Haplodiploidy and paternal genome elimination (HD/PGE) are common in invertebrates, having evolved at least two dozen times, all from male heterogamety (i.e., systems with X chromosomes). However, why X chromosomes are important for the evolution of HD/PGE remains debated. The Haploid Viability Hypothesis posits that X-linked genes promote the evolution of male haploidy by facilitating purging recessive deleterious mutations. The Intragenomic Conflict Hypothesis holds that conflict between genes drives genetic system turnover; under this model, X-linked genes could promote the evolution of male haploidy due to conflicts with autosomes over sex ratios and genetic transmission. We studied lineages where we can distinguish these hypotheses: species with germline PGE that retain an XX/X0 sex determination system (gPGE+X). Because evolving PGE in these cases involves changes in transmission without increases in male hemizygosity, a high degree of X linkage in these systems is predicted by the Intragenomic Conflict Hypothesis but not the Haploid Viability Hypothesis. To quantify the degree of X linkage, we sequenced and compared 7 gPGE+X species’ genomes with 11 related species with typical XX/XY or XX/X0 genetic systems, representing three transitions to gPGE. We find highly increased X linkage in both modern and ancestral genomes of gPGE+X species compared to non-gPGE relatives, and recover a significant positive correlation between percent X linkage and the evolution of gPGE. These are among the first empirical results suggesting a role for intragenomic conflict in the evolution of novel genetic systems like HD/PGE.

**Significance Statement:** Sex determination systems such as haplodiploidy, in which males’ gene transmission is haploid, are surprisingly common, however, the evolutionary paths to these systems are poorly understood. X chromosomes may play a particularly important role, either by increasing survival of males with only maternal genomes, or due to conflicts between X-chromosomal and autosomal genes. We studied X-chromosome gene richness in three arthropod lineages in which males are diploid as adults but only transmit their maternally-inherited haploid genome. We find that species with such atypical systems have far more X chromosomal genes than related diploid species. These results suggest that conflict between genetic elements within the genome drives the evolution of unusual sex determination systems.

## Introduction

Many animal lineages have evolved genetic systems in which females are diploid but males are haploid or effectively haploid, with each male creating genetically identical sperm carrying the single haploid genome originally inherited from his mother (1). Such systems range from haplodiploidy (HD), in which males are produced from unfertilized eggs; to embryonic paternal genome elimination, in which diploid males eliminate their paternal genome early in development; to forms of germline-specific PGE (gPGE), where the paternal genome is present in male diploid cells but excluded during male meiosis (Figure 1a).

**Figure 1.**
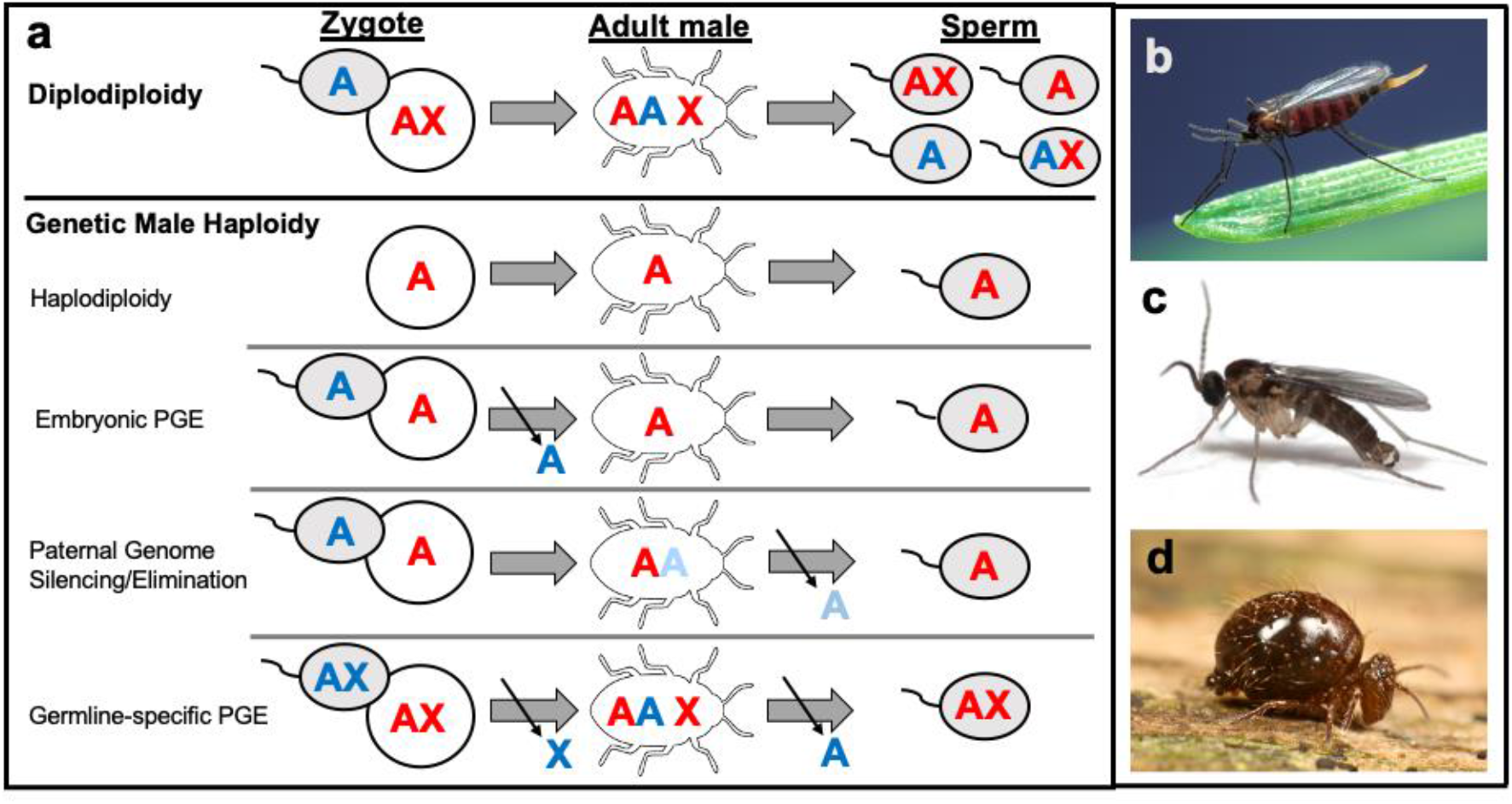
Schematic of different genetic systems discussed. a) Male production and spermatogenesis under diplodiploidy, and various forms of male genetic haploidy are shown. Blue and red letters indicate paternal and maternally derived material respectively. A and X represent autosomes and X chromosomes, respectively. Shown are haplodiploidy (HD), where males develop from unfertilized eggs; embryonic paternal genome elimination, where males eliminate their paternally inherited genome early in development; paternal genome silencing/elimination, a form of germline PGE where males silence paternal autosomes in somatic cells (indicated by the light blue “A” in males) and eliminate these chromosomes during meiosis; and germline-specific PGE (gPGE), as observed in Sciarids, Cecidomyiids and Symphypleonan springtails, wherein males are produced by somatic loss of the paternal X chromosome(s), and the paternal genome is eliminated in spermatogenesis. b-d) representative species with gPGE: b) the Hessian fly gall midge *Mayetiola destructor* (Cecidiomyiidae), image by Scott Bauer and publicly available via the U.S. Department of Agriculture; c) the fungus gnat *Bradysia coprophila* (= *B. tilicola;* Sciaridae), image by Mike Palmer, used with permission; and d) the springtail *Allacma fusca* (Sminthuridae), image publicly available and taken by Andy Murray.

HD/PGE is widespread, seen in ~12% of arthropods and having evolved roughly two dozen times (1). This recurrent evolution perhaps reflects the various advantages of HD/PGE, particularly to mothers, who can increase the transmission of their genes over paternally-inherited genes, control the sex ratio, ensure reproductive success without a mate (in HD), and, under monogamy, reduce conflict between gregarious offspring (2–6). Given these general benefits, why does HD/PGE evolve in some lineages and not in others? An important clue comes from the finding that HD/PGE evolves from ancestral male heterogamety (XX/XY or XX/X0) (7, 8). The most influential hypothesis for this association is the Haploid Viability hypothesis. This hypothesis emphasizes that, starting from an ancestral standard diploid system, newly-evolved haploid males are expected to have markedly lowered fitness due to uncovered recessive deleterious mutations. However, because hemizygosity of X-linked genes facilitates purging of recessive deleterious mutations, an ancestral increase in the proportion of genes on the X chromosome is expected to lead to a decrease in the total number of segregating recessive deleterious mutations, reducing the fitness burden of deleterious mutations for newly-evolved haploid males (5, 8–10).

However other hypotheses are possible. In particular, the Intragenomic Conflict hypothesis, instead, is a more general hypothesis that sees conflicts between genes within individuals as forces that can destabilize genetic systems and thus promote the origins of novel systems including, but not limited to, HD/PGE (11–13). X chromosomes seem to be more often associated with intragenomic conflict compared to autosomes (14–16). In particular, X-linked genes can evolve X chromosome drive (>50% transmission of the X in sperm), which can lead to female-biased population sex ratios. Under such sex biases, males have higher average fitness, thus driving selection for new means of producing males (17, 18). This generally increased male fitness could select for production of haploid males. Moreover, silencing or foregoing the paternal genomic contribution (and in particular the paternal X) could be selectively advantageous insofar as selfish driving X alleles are expected to disproportionately act in males. Although more theoretical work is needed, according to the most developed model, HD/PGE in particular could evolve under the Intragenomic Conflict hypothesis through the exploitation of X chromosome drive by maternal autosomes that increase their transmission by becoming effectively X-linked (11). According to this model, the more genes are X-linked, the more genes will be selected to promote X chromosome drive (and the fewer will be selected to suppress drive), increasing the chance of the evolution of male haploidy.

These two hypotheses differ in whether they predict an association between X linkage and the origins of gPGE in those gPGE systems in which paternal chromosomes are expressed in the soma but are eliminated during male meiosis (Fig. 1a). Given diploid expression of autosomes in the male soma, gPGE, unlike other types of HD/PGE, does not uncover deleterious recessive alleles. Thus, the Haploid Viability hypothesis does not predict an association between X linkage and the evolution of gPGE. However, the notion that X-autosome conflict drives novel systems equally applies to gPGE and other HD/PGE systems, thus the Intragenomic Conflict hypothesis predicts an association between X linkage and the evolution of gPGE. (Notably, in most characterized gPGE systems including those studied here, the paternal genome remains present and expressed through the diploid pre-meiotic stages of spermatogenesis and is only eliminated during meiosis.)

To our knowledge, this differential prediction has not been noted or tested. gPGE systems that retain sex chromosomes and diploid expression of somatic autosomes are known from three lineages: flies in Sciaridae (19) and Cecidomyiidae (20) (fungus gnats and gall midges respectively, two families in the diverse dipteran superfamily Sciaroidea) and springtails in the order Symphypleona (21). Sciaridae and Cecidomyiidae represent a substantial fraction of worldwide biodiversity and are some of the most abundant species of flying insects found in tropical rainforests and in temperate ecosystems, with many new species in these groups continuing to be described (22–24). Sciaridae, Cecidomyiidae, and Symphypleona have independently evolved similar variants of gPGE, in which males are produced through somatic elimination of paternal X chromosomes, while the remainder of the paternal genome is retained until its elimination during meiosis (Fig. 1) (7, 14, 21, 25–30). These clades offer a powerful opportunity to disentangle whether the origin of HD/PGE is better explained by the Haploid Viability hypothesis or the Intragenomic Conflict hypothesis.

To test these two hypotheses for the origins of HD/PGE, we performed whole genome sequencing and comparative analysis of 17 genomes from species with gPGE and their non-gPGE relatives. We developed methods to estimate genome-wide X chromosomal linkage using additional 35 dipteran species for validation, and then used these methods to estimate X linkage across the 17 studied species. We find evidence for ancestral gene-rich X chromosomes coincident with three independent origins of gPGE. These results provide the first empirical evidence for a role for intragenomic conflict in the origins of atypical genetic systems.

## Results and Discussion

### Development and testing of an improved method to estimate genome-wide X chromosomal linkage

Illumina genome sequencing and assembly was performed for males of each studied Sciaroidea species, and average read coverage was calculated for each contig. For the dipteran species, putative orthologs of *D. melanogaster* genes were identified via TBLASTN searches of each genome. Each ortholog was then assigned to one of the so-called Muller elements, *D. melanogaster* chromosomal linkage groups that have persisted over long evolutionary time in some fly lineages (31, 32) (though not all (33)). For each Muller group in each species, the fraction of X-linked genes was estimated from read coverage distributions using improved methods based on Vicoso and Bachtrog’s previous work demonstrating the lability of sex-linked Muller elements across Diptera (33) (see Supplemental Information for full detail). To validate our method, we ran our assignment on 35 dipterans outside of Sciaroidea with a variety of DNA coverage distributions, recapitulating and expanding what was found about Muller element linkage across Diptera by Vicoso and Bachtrog (Fig. S5, Table S1) (33). Publicly available chromosome level assemblies for *B. coprophila, A. gambiae, T. dalmanni*, and several Drosophilid species allowed for direct comparison to our assignment and for each we found our estimation of X linkage to be within 1% of previous estimates, allowing us to be confident in our assignment of X-linked and autosomal genes (Fig. S5a).

### Increased numbers of X-linked genes in gPGE species relative to related species

To test whether the evolution of gPGE is associated with gene-rich X chromosomes, we estimated the proportion of the genome that is X-linked for 17 species of Sciaroidea flies and two species of springtails. We sampled the flies across seven families spanning the root of Sciaroidea, including two families with gPGE and two outgroup species within Bibionomorpha. We used the publicly available genome assembly and annotation supported by physical mapping for the Hessian fly, Cecidomyiid *Mayetiola destructor* (34, 35), and also used available female read data to estimate relative male to female coverage. For the springtails, we performed genomic sequencing of males and females from one species from the gPGE order Symphypleona, *Allacma fusca* (Fig. 1d), and of *Orchesella cincta*, from Entomobryidae, the closest relative springtail order with standard XX/X0 sex determination. In Springtails, instead of orthologs, we used genome annotations to estimate the gene density and used both male and female read coverage data. Our assignment methods provided clear estimates of X linkage for nearly all our species, with exceptions in one species, the Cecidomyiid *Lestremia cinerea*, which showed three distinct peaks in genome coverage rather than two, as well as a low number of complete BUSCO genes present (Fig. S1, S2).

Among all non-gPGE fly species of Bibionomorpha, we found very few X-linked genes, with the X chromosome in all species comprised mostly of genes from the diminutive F Muller element (<1% of all genes), consistent with the previous inference for the ancestral dipteran X chromosome (Fig. 2a) (33). Interestingly, no Muller elements exhibited clear X-linked peaks in coverage in *Platyura marginata* and *Symmerus nobilis*, the latter of which is sister to all other Sciaroidean species, suggesting either homomorphic sex chromosomes, a lack of an X chromosome, or neo-X chromosomes too recently evolved to be distinguished via coverage as previously observed (33).

**Figure 2.**
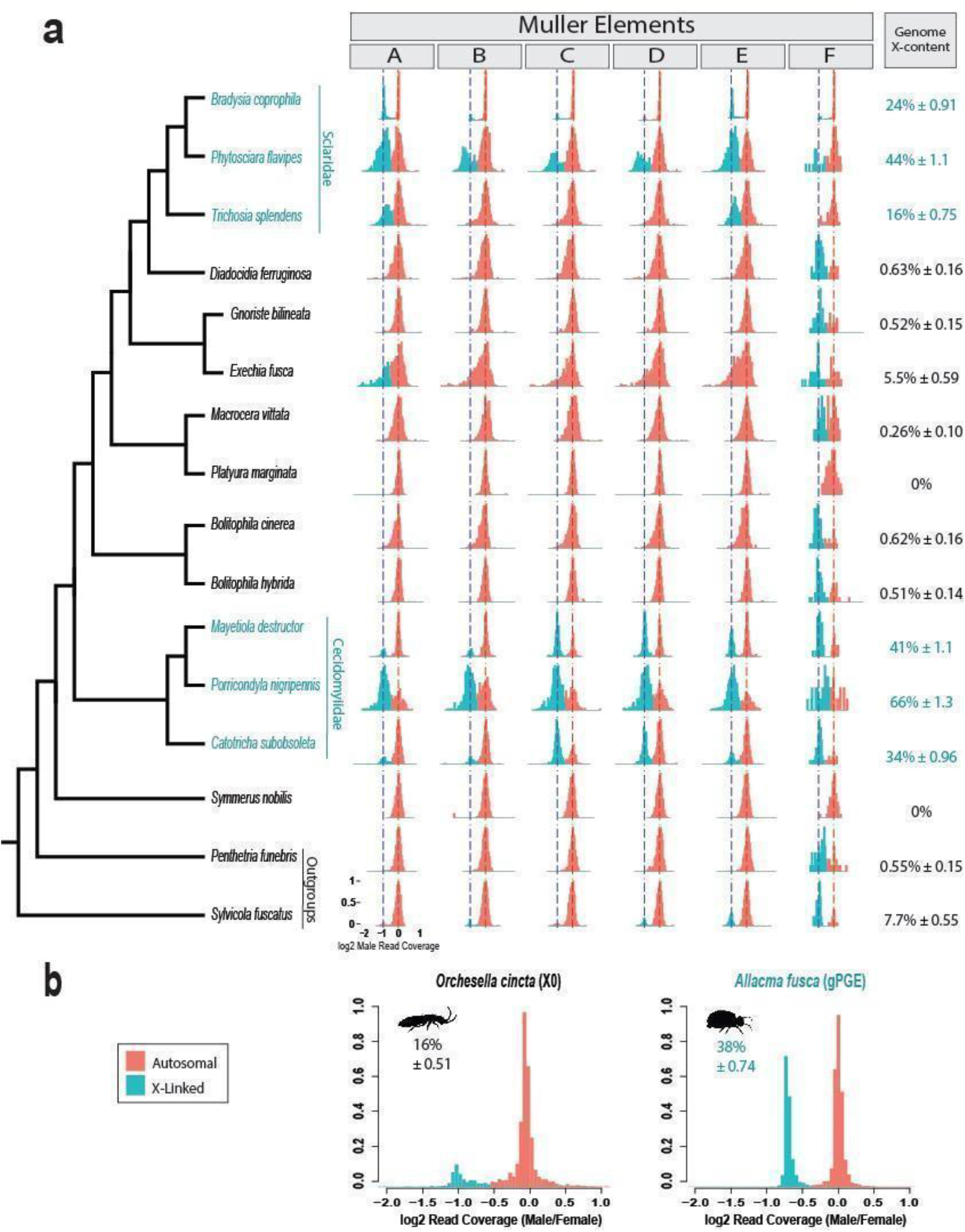
Frequency of X-linked and autosomal genes in gPGE species and related diplodiploid species, assessed by DNA read coverage. a) Sciaroidea and outgroups within Bibionormorpha; topology based on Ševčík et al. (51). Plots for each Muller element show log2 male read coverage normalized by putative median autosomal coverage, with assigned X linkage (blue) and autosomal linkage (red) indicated. The y-axis represents gene frequency scaled to the maximum in each distribution. Red dashed vertical lines at 0 indicate the expected autosomal coverage peak, blue dashed lines at −1 indicate the expected position of the X-linked peak, at half the coverage of the autosomes. Blue and black species names and genome-wide estimates represent gPGE and diplodiploid species, respectively. Percent estimates represent percent X linkage for each Muller and across each full genome, with error represented by 2SD. As with the two springtails, for the Cecidomyiid *Mayetiola destructor*, female read data was available and thus male/female coverage is shown. In *M. destructor* genes are only included if assignments agree with previous physical mapping placements or were previously unassigned (35). b) Whole genome autosomal and X linkage distributions for springtails diplodiploid *Orchesella cincta* and gPGE *Allacma fusca* showing relative male/female coverage.

By contrast, for all six studied gPGE species in both the Sciaridae and Cecidomyiidae clades, genome-wide, we found large fractions of genes to be X-linked, including genes from all six Muller elements (Fig. 2a, 3). Notably, our results agree with previous results for *M. destructor*, identifying Muller elements C, D, F, and E as partially X-linked (33), and our methods additionally detect a small minority of X-linked genes for elements A and B. We also found a clear contrast between the two studied springtail genes: while only 16% of genes in the genome of non-gPGE *Orchesella cincta* are X-linked, for the gPGE springtail *Allacma fusca*, 38% of annotated genes are X-linked (Fig. 2b).

**Figure 3.**
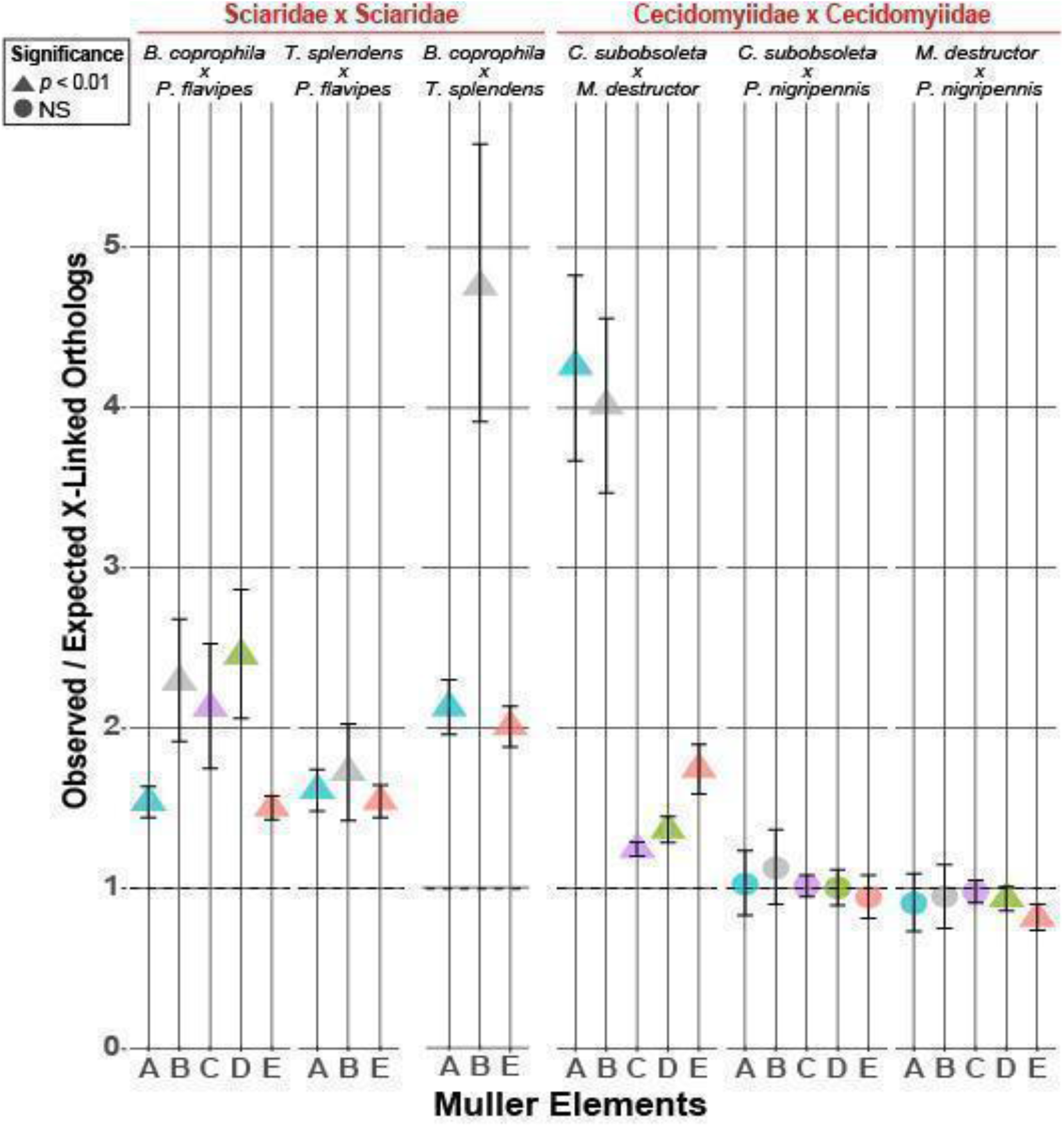
Number of ortholog pairs in which both genes are X-linked, compared to the null expectation, for pairs of gPGE species from the same family. Within-family comparisons are shown, between-family comparisons in Fig. S3. Color indicates Muller element. Muller elements for which species do not share X-linked orthologs are excluded, as is the F element. Shapes indicate significance via Chi square. Error bars represent 95% CIs computed from 10,000 bootstrap replicates. Observed/Expected value if no association between X-linked orthologs is 1.

### Statistical tests support the relationship between gPGE and X linkage

To test the association between percent X chromosome linkage and the evolution of gPGE, we used multiple statistical methods. While the number of transitions to gPGE with X chromosomes is small and all current phylogenetic methods with a binary response variable are prone to inflated Type 1 error rates with small sample sizes (36), we are only aware of those transitions represented here and took several approaches to mitigate this issue. First, we used two different phylogenetically informed methods, a Bayesian probit model and a likelihood model using logistic regression, which gave similar results. In addition, we performed stringent diagnostics of our models, including bootstrapping, checks for Markov chain convergence, and autocorrelation of samples in the posterior distribution.

Our non-phylogenetically informed generalized linear model (glm) inferred a positive and significant effect of the degree of X linkage on the evolution of gPGE, with every one standard deviation increase in percent X linkage increasing the log odds of evolving gPGE by a factor of 3.34. However, occurrences of gPGE group in two tight clusters on the phylogeny, and we detected a strong phylogenetic signal in the amount of X-linkage (Blomberg’s K = 0.946; p = 0.012), meaning this correlation could largely result from phylogenetic proximity. Therefore, we used Bayesian estimation with a mixed phylogenetic model and found a robust, positive effect that does not overlap zero (mean = 1.87, 95% 95% CI = 0.366 - 3.73, pMCMC = 0.005; MCMC effective sample size = 304,300). In addition, we performed Ives and Garland’s binary phylogenetic logistic regression (37) with 10,000 bootstrap replicates and found an even stronger standardized effect (mean = 3.37, 95% bootstrap CI = 3.36 - 3.40, p-value = 2.382e-08) which is similar to the estimate with the non-phylogenetic glm. Thus, all three methods support transitions to gPGE being more prevalent in lineages with a higher proportion of X-linked genes (Fig. S3; see Supplemental Information for full details).

### Correspondence between X-linked genes within families indicates ancestrally gene-rich X chromosomes

Although we found an association between gene-rich X chromosomes and gPGE in all three independent origins of this genetic system, the observed association could be explained by either X linkage facilitating the evolution of gPGE or vice versa. Consistent with the former, we see the same patterns of Muller group X linkage within families (E>A>B in Sciaridae species; C>D>E>A>B in Cecidomyiidae). In addition, we found an association between X-linked gene subsets within individual Muller elements, as expected from ancestral linkage. For instance, the subsets of Muller B genes that are X-linked in the Sciaridae species *B. coprophila* and *T. splendens* significantly overlap, and the same is true for all partially X-linked Mullers in both Sciaridae (Fig. 3). By contrast, X-linked genes between Sciaridae and Cecidomyiidae do not significantly overlap, supporting independent origins of the large X in these two families (Fig. S4).

Examination of Cecidomyiidae reveals an intriguing pattern. The deeply-diverged species *C. subobsoleta* and *M. destructor* show high correspondence between X-linked gene subsets, indicating substantial ancestral X linkage. However, *P. nigripennis* shows divergent X linkage, with no significant pattern seen in shared X linkage with other Cecidomyiids, and a relative increase in X linkage on Muller elements A and B. This pattern suggests turnover and increases in X linkage in this lineage since the divergence from *M. destructor* (or, less parsimoniously, parallel loss of A/B linkage in the other lineages) (Fig. 2a, 3).

### X Chromosome lability and partial Muller linkage

Our data attest to substantial dynamism of the X chromosome and Muller linkage in both gPGE families within Diptera. This is in contrast to the dominant model of Dipteran sex chromosome evolution where sex linked Muller elements are expected to remain stable over long evolutionary periods. Some notable cases indicate remarkable conservation, such as the X chromosome of the German cockroach which has remained conserved with the ancestral dipteran X chromosome (Muller element F) despite 400 million years of divergence (38). On the other hand, even within Drosophila this pattern is disrupted, with fusions of ancestral drosophilid X-linked element A and typically autosomal element D in the obscura clade into the X chromosome, as well as in *D. willistoni* (Fig. 4b) (39, 40). Vicoso and Bachtrog demonstrated abundant sex chromosome turnover across Diptera, broadly challenging the established paradigm of sex linked Muller element stability (33).

In addition to demonstrating cases of lost and replaced sex chromosomes, Vicoso and Bachtrog also showed cases of partial linkage, where parts of multiple Muller elements are incorporated into sex chromosomes (33). Specifically, they find partial linkage for the B element of *Holcocephala fusca* and for the E element of *M. destructor*, both of which our methods also identified as partially X-linked (35% and 40% of genes, respectively). We additionally find minor partial linkage of Muller elements A and B in *M. destructor* (Fig. 2a and Supp figure 4b). In *Anopheles gambiae*, element A is typically discussed as if fully X-linked, however the X chromosome has been previously shown to be only partially composed of element A and parts of other Muller elements, while the rest of ancestrally Muller A genes are now found on autosomes (41), consistent with our results (Fig. S5, Supplemental table 1). Additionally, minor partial X linkage of *A. gambiae* elements E (11%) and F (33%) has been previously identified (42) and is consistent with our findings of 11% and 29% X linkage respectively (Table S1). Our methods demonstrate the resolution to detect low levels of X linkage and suggest partial linkage and general Muller element breakdown may be more common than is generally appreciated.

### Concluding remarks

We find that species in the gPGE groups Cecidomyiidae and Sciaridae have, on average, X chromosomes 37 times more gene-rich than non-gPGE Sciaroidea species, with a more than doubling of the X chromosome gene content of the gPGE springtail species compared to the diploid outgroup (Fig. 2a and b). Furthermore, we recovered a robust positive correlation between the percent X linkage in the genome and the evolution of gPGE (Fig. S3). While having additional independent origins of X chromosome-containing gPGE would add strength to our conclusions, we are only aware of those studied here and our results are bolstered by multiple statistical methods.

Notably, while previous similar reports of an association between the extent of X linkage and atypical sex determination are consistent with either the Haploid Viability hypothesis or the Intragenomic Conflict hypothesis (8), these findings represent the first empirical evidence that suggests Intragenomic Conflict as a strong driver of the evolution of unconventional sex determining systems such as gPGE and haplodiploidy. Given the widespread and repeated evolution of male haploidy, and its association with many unique ecological and life history strategies, our findings point to an important role for intragenomic conflict in shaping biology at all levels from molecule to organism to community.

## Materials and Methods

### Specimens and sequencing

In order to compare X chromosomes of gPGE species to their diplodiploid relatives, we collected and sequenced males of 18 species, 14 belonging to the superfamily Sciaroidea spanning nearly all families within, two outgroup species in the dipteran families Anisopodidae and Bibionidae (Sciaroidea and these families are both in the infraorder Bibionomorpha), and two springtail species, *Allacma fusca* and *Orchesella cincta*. Eleven Bibionomorphan specimens were collected and provided by Jan Ševčík, *Catotricha subobsoleta* by Scott Fitzgerald, *Bolitophila hybrida* by Nikola Burdíková, and *Bradysia coprophila* (= *B. tilicola*) was cultured at the University of Edinburgh. Springtails were provided by Jacintha Ellers. Both specimens were flash-frozen and stored at −80°C.

For 15 dipteran species, DNA extractions (Qiagen DNAeasy Blood & Tissue kit), library preparation (Illumina TruSeq kit), and sequencing (Illumina Hi-Seq) were performed by Iridian Genomes. Genomes were assembled using Megahit 1.13 (43). For the two springtail species and the Sciarid *Bradysia coprophila*, DNA was extracted from male heads using a modified extraction protocol from DNAeasy Blood & Tissue kit (Qiagen, The Netherlands) and Wizard Genomic DNA Purification kit (Promega). TruSeq DNA Nano gel free libraries (350 bp insert) were generated by Edinburgh Genomics (UK) and sequenced on the Illumina HiSeq X (for springtails) or NovaSeq S1 (for *B. coprophila*) generating short reads (150 bp paired-end). The genome for *B. coprophila* was assembled using Megahit 1.2.9 (43). The genome of springtail *A. fusca* was assembled using SPAdes v3.13.1 (44). Both genomes of *B. coprophila* and *A. fusca* assemblies were decontaminated with BlobTools (45). The assembly of *A. fusca* was annotated using BRAKER 2.1.5 (46). We assessed the quality of all genomes using BUSCO (47), to determine the proportion of single copy orthologs expected to be present in either insects (insecta_odb10 for fungus gnat species) or arthropods (for springtails) in the genome assemblies (Fig. S1). *Lestremia cinerea* was excluded from downstream analysis due to irregular genome coverage patterns and a low number of complete BUSCO genes present, indicating likely issues with the genome quality for this species (Fig. S1 and S2). We used publicly available genome assemblies for the Cecidomyiid *Mayetiola destructor* (GCA_000149195.1) and for the springtail *Orchesella cincta* (GCA_001718145.1). For *M. destructor*, we used publicly available male (SRR1738190) and female reads (SRR1738189), and for *O. cincta*, we additionally used available female reads (SRR2222657).

### Assigning ancestral linkage groups

The X chromosome in each fly species was identified using two strategies— Muller group linkage and genomic read coverage, similar to strategies implemented in Vicoso and Bachtrog 2015 (33). Muller elements are six chromosomal elements first characterized in *Drosophila* that are regarded as being informative about chromosomal linkage (31). The *D. melanogaster* proteome (flybase r6.32) (48) was searched against each assessed genome translated into 6 frames using TBLASTN. Top hits for each *D. melanogaster* gene were identified and corresponding genes were classified by the Muller element of their closest *D. melanogaster* ortholog. The X chromosomes in springtails were identified using the coverage approach only.

### Identifying X linkage via coverage

Our second strategy implemented DNA coverage levels to characterize autosomal and X-linked sequence, as we expect the single copy X chromosome in males to cause X-linked sequence to be found at half the coverage level of autosomes. Male DNA reads were mapped to their respective genome assemblies and repetitive sequence that could not be singly mapped was accounted for when calculating an adjusted coverage (See Supplemental methods). For species in which female read data was available, *M. destructor* and the two springtails, the relative coverage of male to female was used. In the case of *A. fusca*, we used median coverage of two males and 11 females available (26). To classify genes by coverage as either autosomal or X-linked, we used a multi-step protocol relying on the full genome and per-Muller male DNA coverage distributions (See detail in Supplemental Information). We also assessed 35 other dipteran genomes outside Bibionormorpha using publicly available data and the same methods of analysis (Fig. S5, Table S1).

### Statistical analysis and phylogenetic correction

To test the association between X linkage and the evolution of PGE, we estimated a Bayesian generalized linear mixed “threshold” model (49) and a likelihood-based phylogenetic logistic regression described in (50) Both methods attempt to control for the phylogenetic relatedness of the species. For full detail, see Supplemental Information.

### Testing for ancestral Muller group linkage

To test for evidence of ancestral X linkage, we compared various pairs of species. We studied each Muller element for which both compared species had partial X linkage, in which the ancestral linkage groups have broken up and are now partially X-linked and partially autosomal. Genomes of each species pair were reciprocally blasted to defined putative pairwise orthologs using TBLASTX. Only best reciprocal hits and orthologs that blasted to the same *D. melanogaster* gene were included in further analysis. Each ortholog pair was then assigned based on its inferred X/autosomal linkage for both species (X-linked/X-linked, X-linked/autosomal, autosomal/X-linked, or autosomal/autosomal). Association between X linkage across between-species orthologs was tested by a Chi square test.

## Acknowledgements and funding sources

We would like to acknowledge Scott Fitzgerald, Nikola Burdíková, Jacintha Ellers for their work collecting specimens. Genome sequencing of all dipteran specimens besides *B. coprophila* was funded by Iridian Genomes. NA and BW were supported by NSF Award #1616878 to SWR. Springtail sequencing and KSJ were supported by a European Research Council Starting Grant (PGErepro, to LR). CH was supported by the National Sciences and Engineering Research Council of Canada postgraduate scholarship and the Darwin Trust of Edinburgh. Springtail collection and LR were supported by a Natural Environment Research Council Independent Research Fellowship (NE/K009516/1) and Dorothy Hodgkin Fellowship (DHF\R1\180120).

## Notes

### Competing Interest Statement

The authors have declared no competing interest.

### Summary of Updates

Added new statistical method (Fig S3; yellow line) and revised text to explain caveats of methods and sample size limitations

