## Supplemental figures and methods for "Gene-rich X chromosomes implicate intragenomic conflict in the evolution of bizarre genetic systems"

Supplemental material

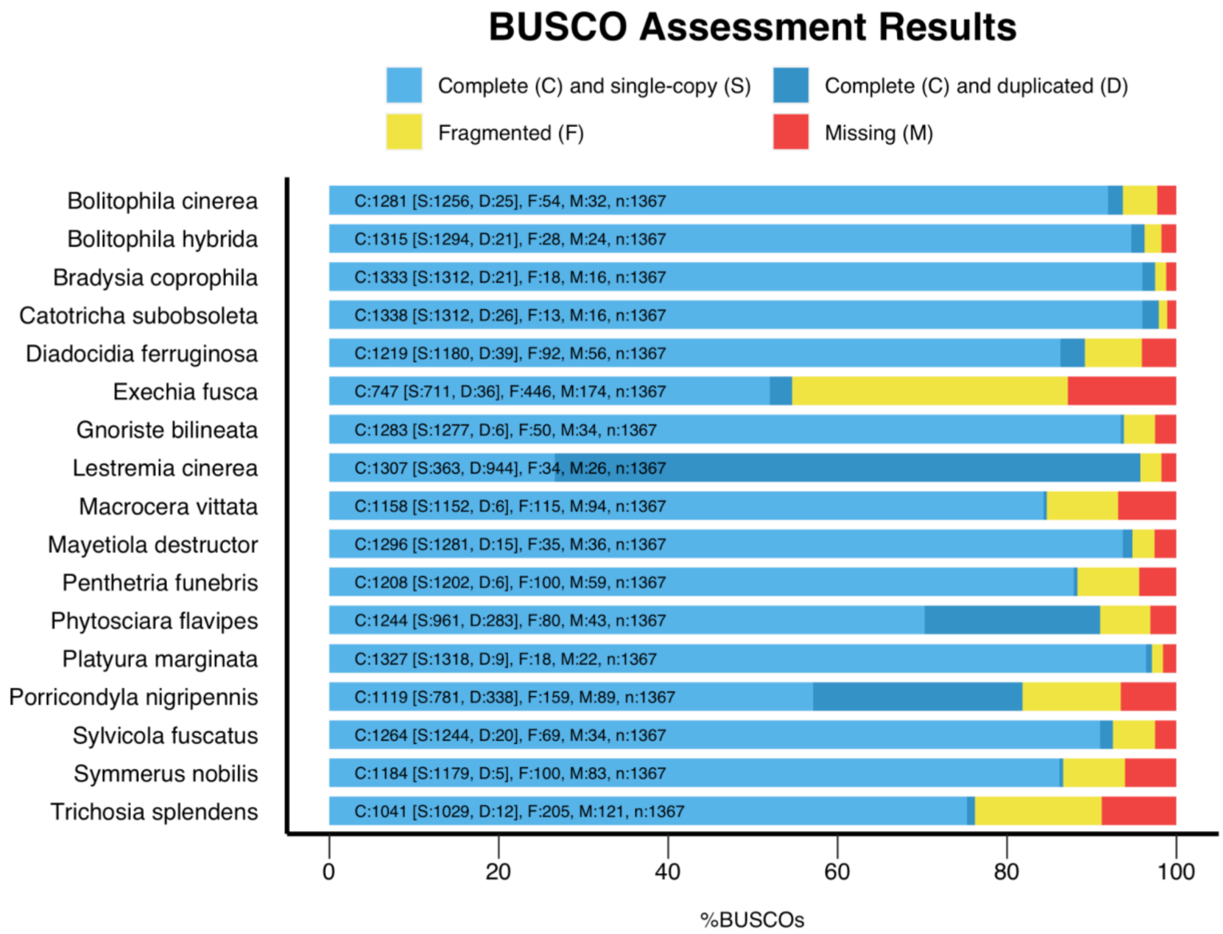

Fig. S1. Genome assembly completeness by BUSCO analysis. Counts for each BUSCO category are shown with abbreviations in the bars.

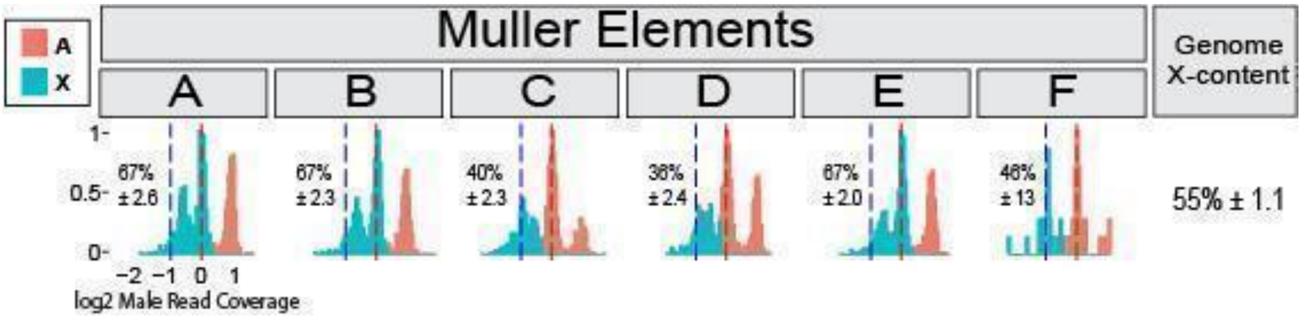

Fig. S2. Male DNA coverage distributions for the Cecidomyid *Lestremia cinerea*. Three distinct read coverage peaks were found in *L. cinerea*, present across all Muller elements. Our method is not designed to accommodate such multi-peak situations, and unsurprisingly breaks down in *L. cinerea*. Our current methods assign a majority of the *L. cinerea* genome as X-linked (55%), though because of the unknown identity of the far left peak, this estimate may be unreliable. This unusual distribution could be indicative of partial genome duplication, as suggested by the BUSCO results, but more investigation is needed.

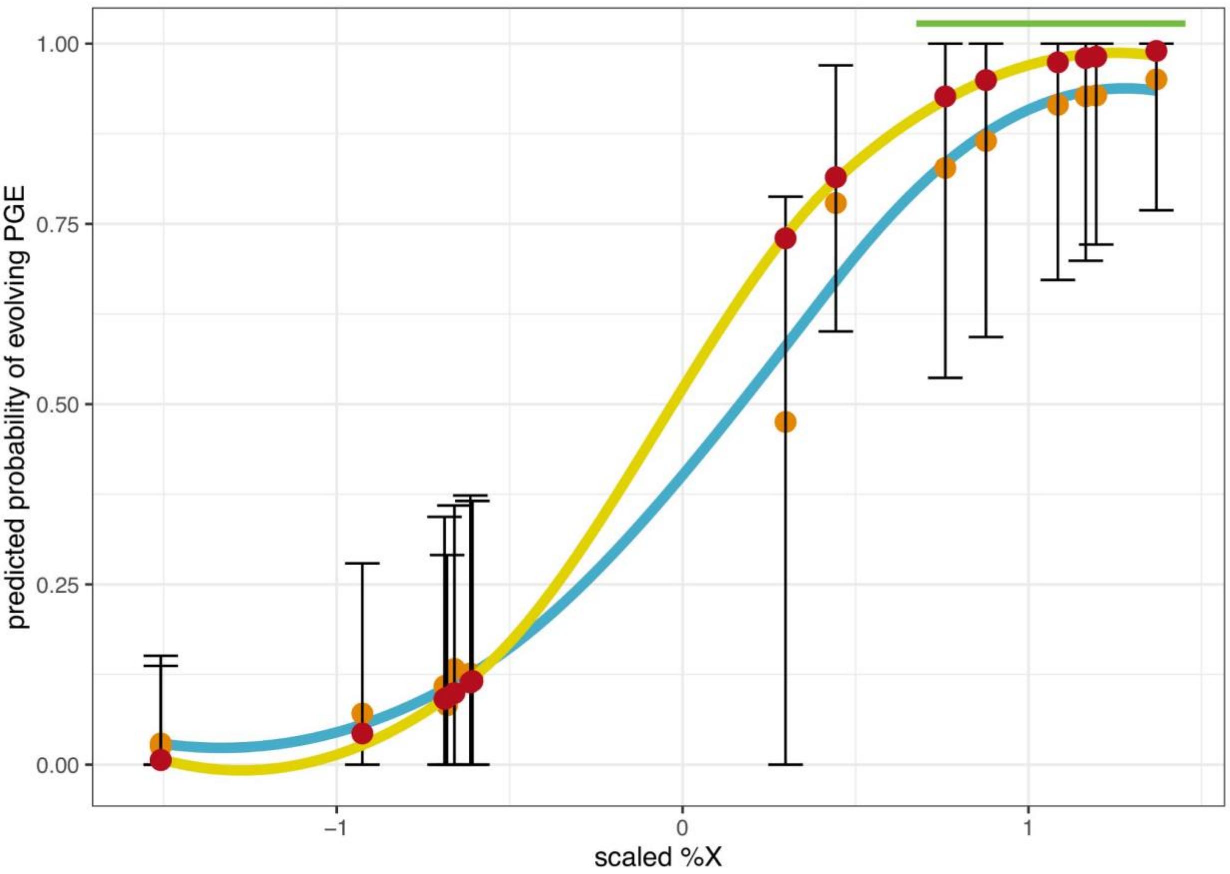

Fig. S3. Predicted probability of evolving paternal genome elimination (gPGE) based on percent X linkage, as estimated from the data. Orange dots are the average, out of sample, predicted probability of gPGE with the MCMCglmmRAM threshold model, and black bars are the 95% confidence interval. The blue line is the average predicted probability made with the geom_smooth function in R (method = "loess"). The yellow line is the logistic regression curve made with the fitted coefficients from the phyloglm model using the plogis function in R. Red dots are the predicted fit from the non-phylogenetic GLM, and the green bar spans the six gPGE species.

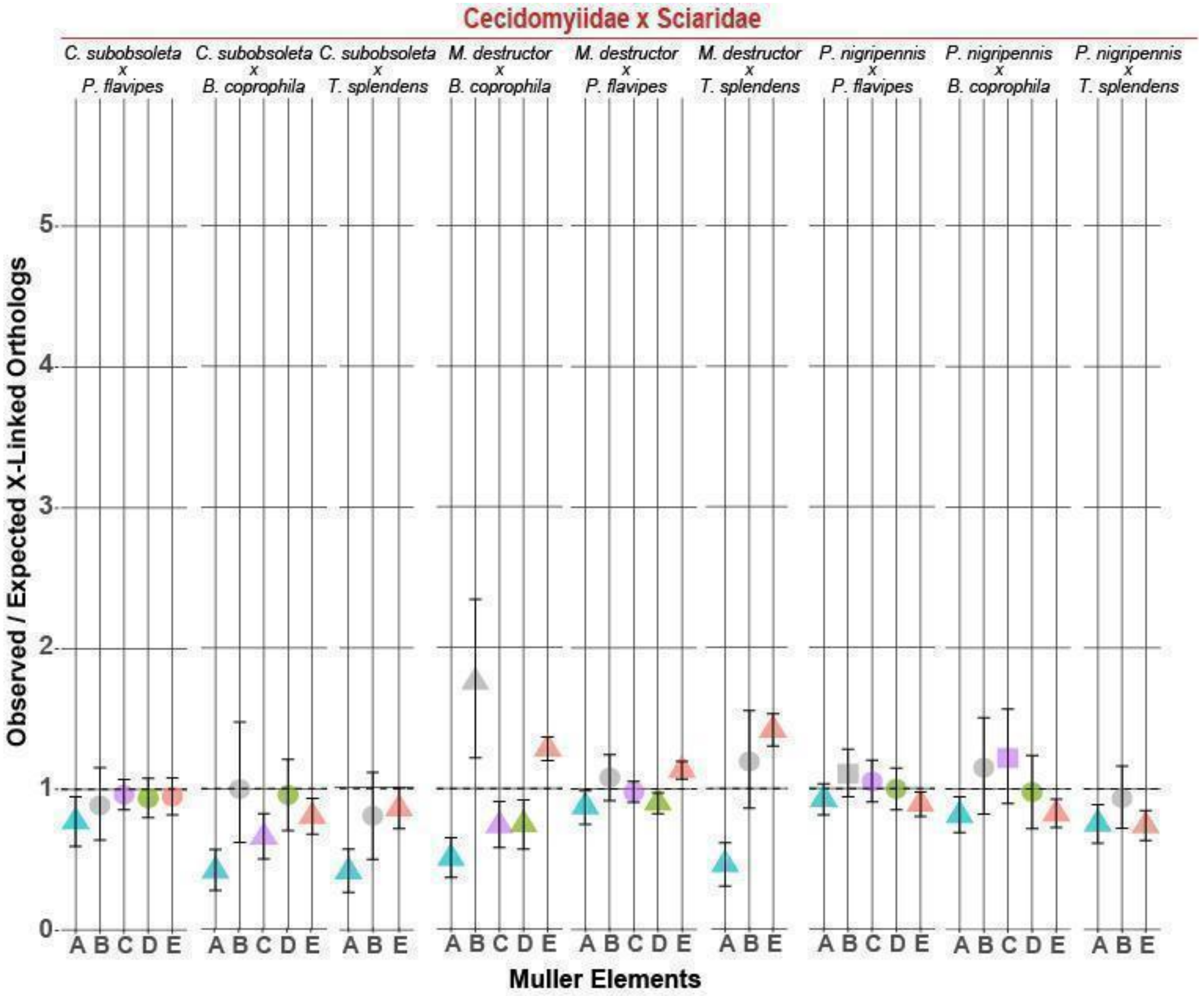

Fig. S4. Number of ortholog pairs in which both genes are X-linked, compared to the null expectation, for pairs of gPGE species from different families. Between-family comparisons are shown here, while within-family comparisons are shown in main Figure 3. Color indicates Muller element. Muller elements for which species do not share X-linked orthologs are excluded, as is the F element. Shapes indicate significance via Chi square. Error bars represent 95% CIs computed from 10,000 bootstrap replicates. Expected value if no association between X-linked orthologs is 1.

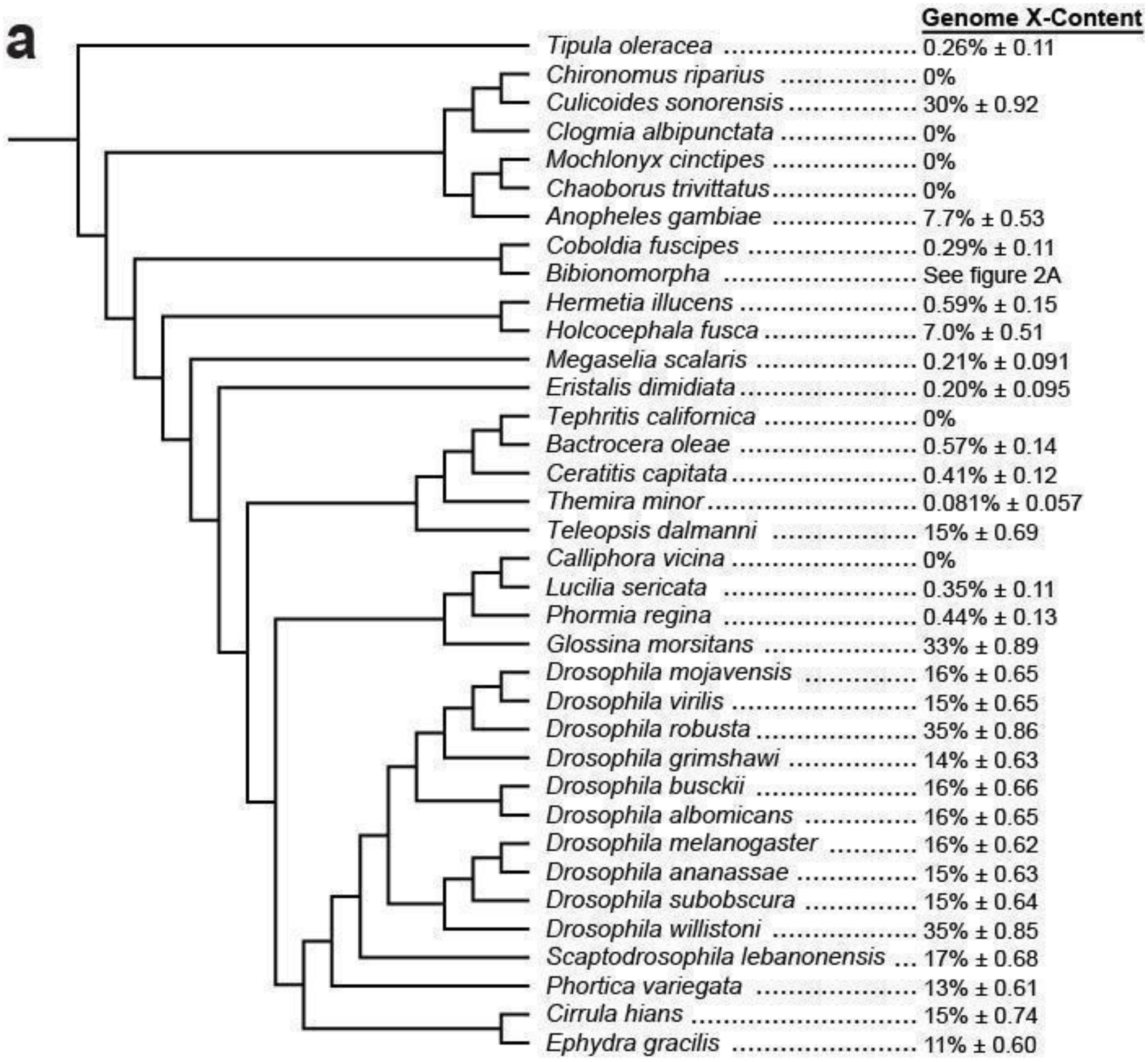
a
Genome X-Content
Tipula oleracea 0.26% ± 0.11
Chironomus riparius 0%
Culicoides sonorensis 30% ± 0.92
Clogmia albipunctata 0%
Mochlonyx cinctipes 0%
Chaoborus trivittatus 0%
Anopheles gambiae 7.7% ± 0.53
Coboldia fuscipes 0.29% ± 0.11
Bibionomorpha See figure 2A
Hermetia illucens 0.59% ± 0.15
Holcocephala fusca 7.0% ± 0.51
Megaselia scalaris 0.21% ± 0.091
Eristalis dimidiata 0.20% ± 0.095
Tephritis californica 0%
Bactrocera oleae 0.57% ± 0.14
Ceratitis capitata 0.41% ± 0.12
Themira minor 0.081% ± 0.057
Teleopsis dalmanni 15% ± 0.69
Calliphora vicina 0%
Lucilia sericata 0.35% ± 0.11
Phormia regina 0.44% ± 0.13
Glossina morsitans 33% ± 0.89
Drosophila mojavensis 16% ± 0.65
Drosophila virilis 15% ± 0.65
Drosophila robusta 35% ± 0.86
Drosophila grimshawi 14% ± 0.63
Drosophila busckii 16% ± 0.66
Drosophila albomicans 16% ± 0.65
Drosophila melanogaster 16% ± 0.62
Drosophila ananassae 15% ± 0.63
Drosophila subobscura 15% ± 0.64
Drosophila willistoni 35% ± 0.85
Scaptodrosophila lebanonensis 17% ± 0.68
Phortica variegata 13% ± 0.61
Cirrula hians 15% ± 0.74
Ephydra gracilis 11% ± 0.60

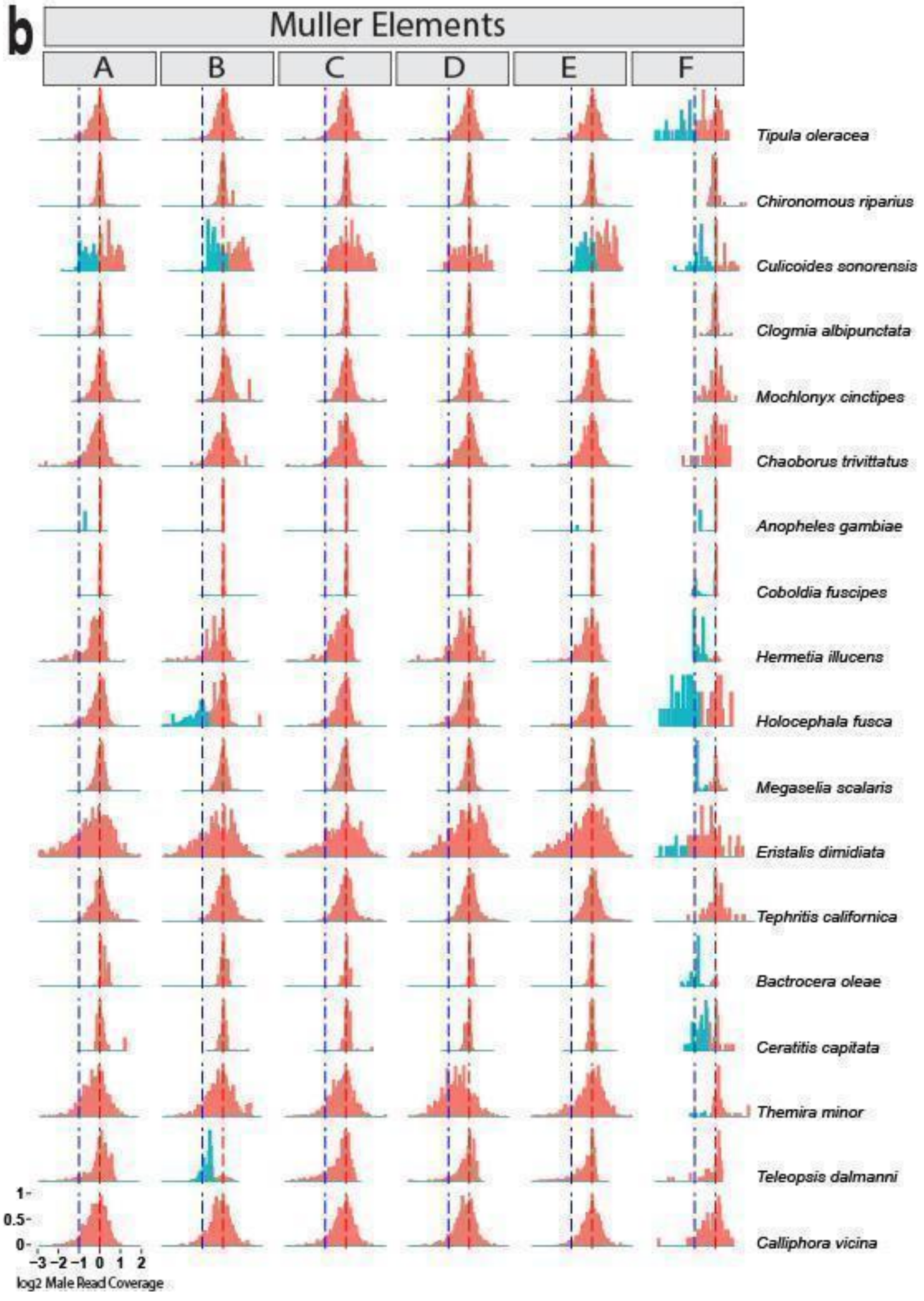
b
Muller Elements
A
B
C
D
E
F
Tipula oleracea
Chironomous riparius
Culicoides sonorensis
Clogmia albipunctata
Mochlonyx cinctipes
Chaoborus trivittatus
Anopheles gambiae
Coboldia fuscipes
Hermetia illucens
Holocephala fusca
Megaselia scalaris
Eristalis dimidiata
Tephritis californica
Bactrocera oleae
Ceratitis capitata
Themira minor
Teleopsis dalmanni
Calliphora vicina
1
0.5
0
−3 −2 −1 0 1 2
log2 Male Read Coverage

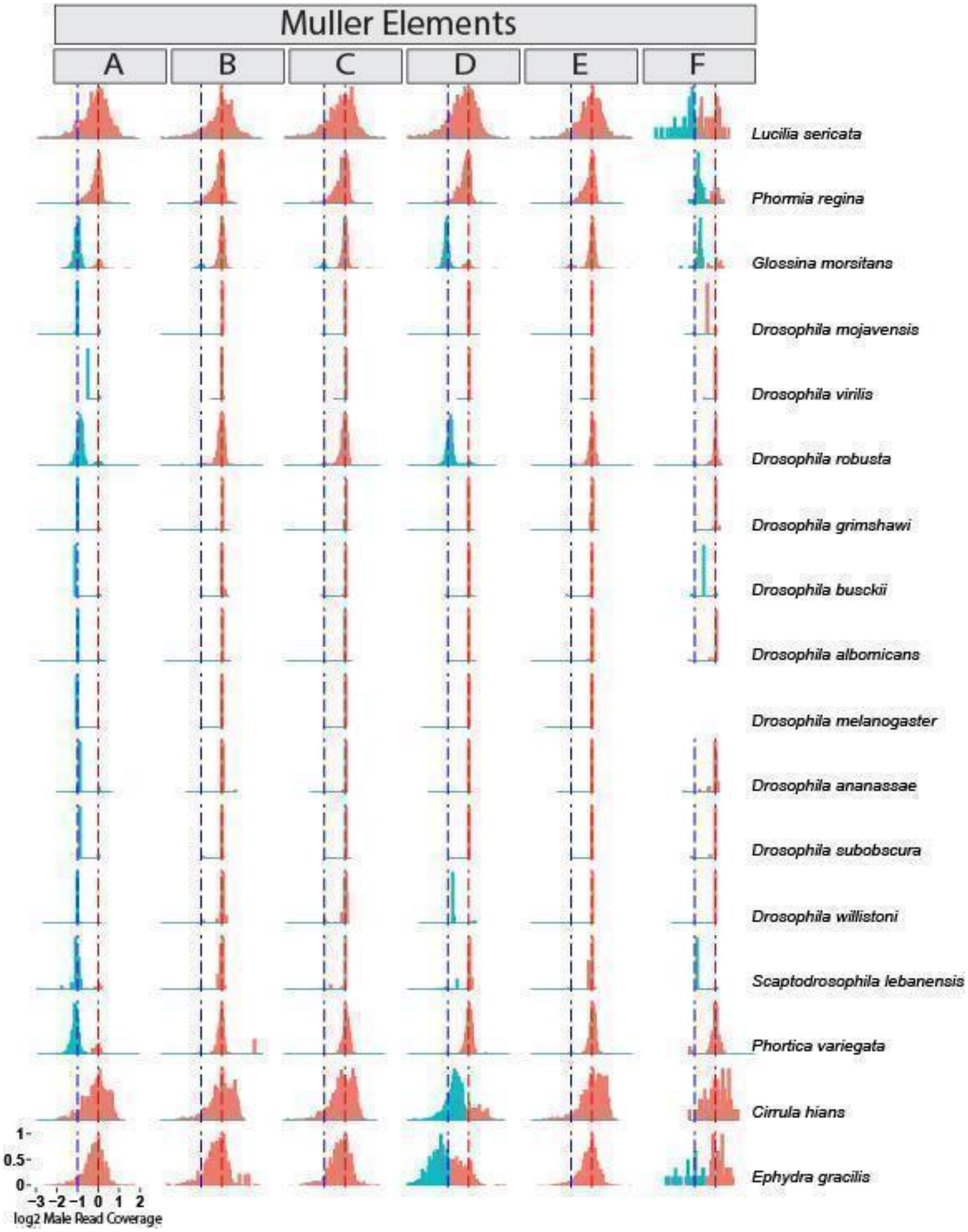

Figure S5. Distribution of X-linked genome content across diverse dipterans, assessed by male DNA read coverage. a) Phylogenetic tree showing the percent of X-linked genes estimated from the whole genome. b) Frequency of X-linked genes across Muller elements, showing log2 male read coverage normalized by putative median autosomal coverage, with assigned X linkage (blue) and autosomal linkage (red) indicated. The y-axis represents gene frequency scaled to the maximum in each distribution. Red dashed vertical lines at 0 indicate the expected autosomal coverage peak, blue dashed lines at -1 indicate the expected position of the X-linked peak, at half the coverage of the autosomes. Percent estimates represent percent X linkage for each Muller and across each full genome, with error represented by 2SD. *Drosophila melanogaster* F element, the dot chromosome, is excluded for insufficient gene number.

Table S1. The estimated percentage of the full genome that is X-linked and the per-Muller element estimates for Bibionormorphan and non-Bibionormorphan dipterans. Error is represented as twice the standard deviation.

| Species | X-linked percent of genome | A | B | C | D | E | F |
|---|---|---|---|---|---|---|---|
| Bibionomorphan species included in main text figures | | | | | | | |
| *Sciara coprophila* | 24% ± 0.91 | 43% ± 2.6 | 13% ± 1.7 | 12% ± 1.6 | 0% | 44% ± 2 | 8.6% ± 6.7 |
| *Phytosciara flavipes* | 44% ± 1.1 | 57% ± 2.6 | 35% ± 2.4 | 33% ± 2.3 | 35% ± 2.3 | 58% ± 2.1 | 25% ± 11 |
| *Trichosia splendens* | 16% ± 0.75 | 38% ± 2.6 | 0% | 0% | 0% | 39% ± 2 | 0% |
| *Diadocidia ferruginosa* | 0.63% ± 0.16 | 0% | 0% | 0% | 0% | 0% | 78% ± 9.6 |
| *Gnoriste bilineata* | 0.52% ± 0.15 | 0% | 0% | 0% | 0% | 0% | 70% ± 11 |
| *Exechia fusca* | 5.5% ± 0.59 | 33% ± 3.2 | 0% | 0% | 0% | 0% | 55% ± 14 |
| *Macrocera vittata* | 0.26% ± 0.1 | 0% | 0% | 0% | 0% | 0% | 46% ± 14 |
| *Platyura marginata* | 0% | 0% | 0% | 0% | 0% | 0% | 0% |
| *Bolitophila cinerea* | 0.62% ± 0.16 | 0% | 0% | 0% | 0% | 0% | 79% ± 9.5 |
| *Bolitophila hybrida* | 0.51% ± 0.14 | 0% | 0% | 0% | 0% | 0% | 68% ± 11 |
| *Mayetiola destructor* | 41% ± 1.1 | 17% ± 2.1 | 17% ± 1.9 | 70% ± 2.3 | 59% ± 2.6 | 40% ± 2.2 | 75% ± 11 |
| *Porricondyla nigripennis* | 66% ± 1.3 | 67% ± 3.1 | 59% ± 3.1 | 68% ± 2.7 | 64% ± 2.9 | 71% ± 2.3 | 66% ± 14 |
| *Catotricha subobsoleta* | 34% ± 0.96 | 17% ± 1.9 | 16% ± 1.8 | 67% ± 2.1 | 42% ± 2.3 | 23% ± 1.7 | 79% ± 9.2 |
| *Lestremia cinerea* | 55% ± 1.1 | 67% ± 2.6 | 67% ± 2.3 | 40% ± 2.3 | 36% ± 2.4 | 67% ± 2 | 46% ± 13 |
| *Symmerus* | 0% | 0% | 0% | 0% | 0% | 0% | 0% |

| | | | | | | | |
|---|---|---|---|---|---|---|---|
| *nobilis* | | | | | | | |
| *Penthetria funebris* | 0.55% ± 0.15 | 0% | 0% | 0% | 0% | 0% | 72% ± 10 |
| *Sylvicola fuscatus* | 7.7% ± 0.55 | 0% | 4.6% ± 1 | 0% | 8.8% ± 1.4 | 17% ± 1.5 | 82% ± 8.8 |
| *Non-Bibionomorphan dipterans* | *X-linked percent of genome* | *A* | *B* | *C* | *D* | *E* | *F* |
| *Tipula oleracea* | 0.26% ± 0.11 | 0% | 0% | 0% | 0% | 0% | 36% ± 12 |
| *Chironomus riparius* | 0% | 0% | 0% | 0% | 0% | 0% | 0% |
| *Culicoides sonorensis* | 30% ± 0.92 | 46% ± 2.5 | 56% ± 2.3 | 0% | 0% | 48% ± 2 | 57% ± 11 |
| *Clogmia albipuntata* | 0% | 0% | 0% | 0% | 0% | 0% | 0% |
| *Mochlonyx cinctipes* | 0% | 0% | 0% | 0% | 0% | 0% | 0% |
| *Chaoborus trivittatus* | 0% | 0% | 0% | 0% | 0% | 0% | 0% |
| *Anopheles gambiae* | 7.7% ± 0.53 | 29% ± 2.3 | 0% | 0% | 0% | 11% ± 1.3 | 29% ± 11 |
| *Coboldia fuscipes* | 0.29% ± 0.11 | 0% | 0% | 0% | 0% | 0% | 39% ± 11 |
| *Hermetia illucens* | 0.59% ± 0.15 | 0% | 0% | 0% | 0% | 0% | 79% ± 9.1 |
| *Holcocephala fusca* | 7.0 % ± 0.51 | 0% | 35% ± 2.2 | 0% | 0% | 0% | 66% ± 11 |
| *Megaselia abdita* | 0.21% ± 0.091 | 0% | 0% | 0% | 0% | 0% | 31% ± 11 |
| *Eristalis dimidiata* | 0.20% ± 0.095 | 0% | 0% | 0% | 0% | 0% | 25% ± 10 |
| *Tephritis californica* | 0% | 0% | 0% | 0% | 0% | 0% | 0% |
| *Bactrocera oleae* | 0.57% ± 0.14 | 0% | 0% | 0% | 0% | 0% | 82% ± 8.7 |
| *Ceratitis capitata* | 0.41% ± | 0% | 0% | 0% | 0% | 0% | 59% ± 11 |

| | | | | | | | |
|---|---|---|---|---|---|---|---|
| | 0.12 | | | | | | |
| *Themira minor* | 0.081% ± 0.057 | 0% | 0% | 0% | 0% | 0% | 10% ± 6.9 |
| *Teleopsis dalmanni* | 15% ± 0.69 | 0% | 81% ± 1.8 | 0% | 0% | 0% | 0% |
| *Calliphora vicina* | 0% | 0% | 0% | 0% | 0% | 0% | 0% |
| *Lucilia sericata* | 0.35% ± 0.11 | 0% | 0% | 0% | 0% | 0% | 51% ± 12 |
| *Phormia regina* | 0.44% ± 0.13 | 0% | 0% | 0% | 0% | 0% | 66% ± 11 |
| *Glossina morsitans* | 33% ± 0.89 | 84% ± 1.8 | 11% ± 1.4 | 10% ± 1.3 | 81% ± 1.7 | 0% | 79% ± 9.4 |
| *Drosophila mojavensis* | 16% ± 0.65 | 100% ± 0 | 0% | 0% | 0% | 0% | 5.1% ± 4.9 |
| *Drosophila virilis* | 15% ± 0.65 | 100% ± 0 | 0% | 0% | 0% | 0% | 0% |
| *Drosophila robusta* | 35% ± 0.86 | 100% ± 0 | 0% | 0% | 100% ± 0 | 0% | 0% |
| *Drosophila grimshawi* | 14% ± 0.63 | 88% ± 1.5 | 0% | 0% | 0% | 0% | 0% |
| *Drosophila busckii* | 16% ± 0.66 | 100% ± 0 | 0% | 0% | 0% | 0% | 100% ± 0 |
| *Drosophila albomicans* | 16% ± 0.65 | 100% ± 0 | 0% | 0% | 0% | 0% | 0% |
| *Drosophila melanogaster* | 16% ± 0.62 | 100% ± 0 | 0% | 0% | 0% | 0% | 0% |
| *Drosophila ananassae* | 15% ± 0.63 | 100% ± 0 | 0% | 0% | 0% | 0% | 0% |
| *Drosophila subobscura* | 15% ± 0.64 | 100% ± 0 | 0% | 0% | 0% | 0% | 0% |
| *Drosophila willistoni* | 35% ± 0.85 | 100% ± 0 | 0% | 0% | 100% ± 0 | 0% | 0% |
| *Scapto-drosophila lebanonensis* | 17% ± 0.68 | 91% ± 1.3 | 0% | 0% | 13% ± 1.4 | 0% | 100% ± 0 |
| *Phortica variegata* | 13% ± 0.61 | 83% ± 1.8 | 0% | 0% | 0% | 0% | 0% |
| *Cirrula hians* | 15% ± 0.74 | 0% | 0% | 0% | 74% ± 2 | 0% | 0% |
| *Ephydra gracilis* | 11% ± 0.60 | 0% | 0% | 0% | 59% ± 2.1 | 0% | 38% ± 11 |

Supplemental information

*Identifying X linkage via coverage*

We used expected DNA coverage levels to distinguish X-linked and autosomal sequence: Because the X chromosome is present in a single copy in males, in males, sequence that is X-linked is expected to be at half coverage compared to autosomal sequence. Trimmed to 50 nucleotides, Male DNA reads for each Bibionomorphan were mapped to their respective genome assemblies using Bowtie with default parameters except for the addition of the -m1 flag to discard reads that mapped to multiple locations in the genome. Because some Bibionomorphan contigs contained large amounts of repetitive sequence that prevented reads from mapping singly, we corrected coverage estimates to only account for singularly mappable positions on the contigs. To do this, we simulated 50nt reads from every mappable position on each contig, mapped them back to the genome from which they were generated using Bowtie, and subtracted the number of reads from each contig that were unable to map singularly from the contig length. This provided us with an adjusted contig length that excluded sequence content that could not be mapped to singularly to use for adjusting coverage estimates; contigs with less than 1000 mappable bases were excluded. Coverage was calculated as: (Read count x read length) / (Contig length - number of multiply mapping reads for that contig + 1). Because male and female DNA sequence for *M. destructor* is available, the comparison of male to female read coverage was used as with the springtails, in addition to using linkage information previously established by physical mapping to more stringently classify X linkage (1)**.**

To assign genes as autosomal or X-linked via coverage, we used a multi-step protocol. First, we used standard methods to (i) identify the highest peak in the full genome coverage distribution and (ii) identify the highest peak in each Muller element distribution that falls closest to the full genome distribution highest peak. The highest peaks were assigned as autosomal centers in all species except *Porricondyla nigripennis* and *Drosophila willistoni,* where the X chromosomal peak was larger.

Next, to detect a second coverage peak, we counted genes in coverage bins outward from the highest peak to the left or right, depending on whether the search for the secondary peak was for the X-chromosomal peak, expected to be at half coverage, or autosomal peak, expected to be twice the X-linked coverage. Each bin count was compared via Chi square to an expected bin count calculated as the average between the bin count and the adjacent bin count, such that only significant rises in coverage like a true peak could be identified. Next, we searched for the second highest peak center within bins with the most significant increase and identified those as potentially X-linked (with the exception of the two species listed above where the secondary peaks were autosomal). Distributions with candidate secondary peaks were labeled as bimodal if the secondary peak was at least one tenth of the height of the highest peak, otherwise unimodal. Genes per Muller element distribution were assigned as X-linked or autosomal via k-means clustering, using the Muller-specific X and autosomal peaks as initial cluster centers. The doubled standard deviation of the proportion of X-linked genes in each distribution was estimated as a proxy for the 95% confidence interval.

*Statistical analysis and phylogenetic correction*

To test the statistical significance of the association between gPGE and the X-linked proportion of the genome within Bibionomorpha, we used Bayesian estimation with a mixed phylogenetic model. The distribution of the predictor variable, the percent of the genome which is assigned as X-linked, includes several zeros and is overdispersed. We therefore transformed it using a Box-Cox power transformation (shifting parameter lambda2 = 0.1; maximum likelihood estimated transformation parameter lambda.hat = -0.1; R package AID v. 2.6 (2)), confirming the normality of the data using a Shapiro-Wilk test ($p = 0.0565$). Before proceeding, percent X linkage was also centered at zero and scaled to improve model specification and ease the interpretation of priors and parameters.

Binomial logistic regression using R's generalized linear model function (glm(gPGE~%X-1, family=binomial(link="logit"))) predicts a positive and significant effect of the degree of X linkage on the evolution of gPGE, with every one increase in the standard deviation of percent X linkage increasing the log odds of evolving gPGE by 3.343. We then used the wald.test function of the aod package v 1.3.1 (3) and calculated a chi-squared test statistic of 5.4, with one degree of freedom, and a p-value of 0.02, indicating that the effect of percent X linkage in the genome is statistically significant on the evolution of gPGE. To see how well this non-phylogenetically informed model performs, we tested whether it fits the data better than a null model (with just an intercept) by calculating the difference in deviance (14.586). Next, we performed a Chi-square test with one degree of freedom to obtain an associated p-value of 0.00013, telling us the model with percent X as a predictor of gPGE fits significantly better than a null model. Finally, we calculated various diagnostic statistics with the R package modEvA v. 2.0 (4), which all showed a strong correlation between percent X linkage and the evolution of gPGE (D2 = 0.658; R2 Cox-Snell = 0.572; R2 Tjur = 0.756; R2 McFadden = 0.641; R2 Nagelkerke = 0.78).

However, this correlation could result from increased genomic conflict in lineages with greater X linkage, as predicted by the Intragenomic Conflict Hypothesis, or simply from shared phylogenetic history. The latter is especially of concern because occurrences of gPGE occur in two tight clusters on the phylogeny. Therefore, to incorporate phylogenetic contrasts, we first obtained a tree with branch lengths based on Ševčík et al. (5). Branch lengths for two species missing from the Ševčík tree, *P. marginata* and *B. hybrida* were calculated as the average of their clades and branch lengths for three other absent species, *Symmerus nobilis, Trichosia splendens, Macrocera vittata,* and *Exechia fusca*, were assigned by proxy as branch lengths from congenitor species in the tree. We then used the chronos function in ape v. 5.5 (6) to obtain time-calibrated branch lengths using penalized likelihood with a "correlated" evolution model. We did this over absolute time by calibrating the rootage to 160 mya, the divergence time for the family Bibionomorpha in the dipteran Timetree of Life (7). For comparison, we also generated a tree with Grafen's (8) branch lengths (power = 1).

We first calculated the phylogenetic signal of the continuous predictor variable, percent X linkage, using Pagel's lambda and Blomberg's K with the phylosig function in the R package phytools v. 0.7-90 (9). Lambda ~ 1 using both the chronogram (1.03; p = 0.010) and the Grafen's branch length tree (0.965; p = 0.009), meaning the evolution of percent X linkage in the genome along the phylogeny roughly corresponds to a Brownian motion expectation. Testing Blomberg's K with our chronogram, percent X linkage in our 16 Bibionomorphan taxa is again as expected under Brownian motion with K = 0.946 (p = 0.012). Using Grafen's branch lengths results in a somewhat lower phylogenetic signal, but still highly significant (K=0.706, p=0.001).

Since percent X linkage and gPGE could be correlated largely due to phylogenetic proximity, we analyzed the evolution of gPGE with a Bayesian generalized linear mixed model using MCMCglmmRAM (10), fitting a reduced version of Wright's threshold model (11–13). The results of this model were generally robust to prior specification but, to improve precision and the efficiency of the MCMC sampling, we used a parameter expanded prior for the variance of the covariance matrix that is strongly skewed toward low variances of phylogenetic signal (prior = list(R = list(V = 1,fix = 1),G = list(G1 = list(V = 1, nu = 1000, alpha.mu = 0, alpha.V = 1)))). We ran the model for 50 million iterations with a burn-in period of 50,000, saving every 100th iteration and using slice sampling to update the latent variables. We inspected MCMC chains visually, and all fixed and random effect variances passed the Heidelberg convergence and half-width diagnostics in the R package coda v. 0.19-4 (14), which tests the null hypothesis that the sampled values come from a stationary distribution. Moreover, autocorrelation was essentially nonexistent for the fixed and random effects variances. The positive correlation between percent X linkage and gPGE described in our results uses our chronogram but is also robust with Grafen's branch lengths (mean = 1.613, 95% CI = 0.2221 - 3.3602, pMCMC = 0.0097;

effective sample size = 195,625). To better understand how our model is performing, we used the predict2 function of the postMCMCglmm package in R v. 0.1-2 (15) to calculate the average marginal, out-of-sample predicted probability of evolving PGE using all of the posterior samples. Unfortunately, post-MCMCglmm only supports ordinal models. Fortunately, the threshold model is identical to an ordinal model, except the residual variance now refers to the variance of the link function rather than the variance of the non-identified residuals. Therefore, we multiplied our threshold model's location effects by sqrt(2) and the variance components by 2, making the predictions equivalent.

Finally, for comparison, we used the phyloglm function of the R package "phylolm" (16) to fit the phylogenetic logistic regression described in Ives and Garland (17) using Firth's penalized likelihood with 10,000 parametric bootstrap replicates. For lower X linkage, the predictions from the three models mostly overlap. However, the Bayesian GLMM threshold model made lower predictions for the evolution of gPGE at high X linkage percentages; in contrast, the phylogenetic logistic regression was in line with the non-phylogenetic GLM (Fig. S3). Nonetheless, all three methods support an association between a higher proportion of X-linked genes and transitions to gPGE.
